# Persistent broken chromosome inheritance drives genome instability

**DOI:** 10.1101/2020.08.26.268565

**Authors:** Chen-Chun Pai, Samuel C. Durley, Wei-Chen Cheng, Nien-Yi Chiang, Boon-Yu Wee, Carol Walker, Stephen E. Kearsey, Francesca Buffa, Johanne M. Murray, Timothy C. Humphrey

## Abstract

Persistent DNA damage arising from unrepaired broken chromosomes or telomere loss can promote DNA damage checkpoint adaptation, and cell cycle progression, thereby increasing cell survival but also genome instability. However, the nature and extent of such instability is unclear. We show, using *Schizosaccharomyces pombe*, that inherited broken chromosomes, arising from failed homologous recombination repair, are subject to cycles of <u>se</u>gregation, DNA <u>r</u>eplication and extensive end-<u>p</u>rocessing, termed here SERPent cycles, by daughter cells, over multiple generations. Following Chk1 loss these post-adaptive cycles continue until extensive processing through inverted repeats promotes annealing, fold-back inversion and a spectrum of chromosomal rearrangements, typically isochromosomes, or chromosome loss, in the resultant population. These findings explain how persistent DNA damage drives widespread genome instability, with implications for punctuated evolution, genetic disease and tumorigenesis.

**One Sentence Summary:** Replication and processing of inherited broken chromosomes drives chromosomal instability.

## Main Text

The DNA damage checkpoint arrests cell division in response to chromosomal breaks thereby facilitating repair (*1*). However, persistent DNA damage resulting from unrepaired DNA double-strand breaks (DSBs) or telomere loss can lead to checkpoint adaptation and cell division (*2-5*). Subsequently, daughter cells inherit unrepaired broken chromosomes leading to genome instability (*6-8*). While chromosomal instability (CIN), a common form of genome instability associated with chromosome loss or alterations, drives intratumoral heterogeneity, copy number variation and drug resistance (*9*), the mechanism by which persistent DNA damage contributes to CIN is poorly understood.

We have previously shown that a DSB induced within a non-essential stable minichromosome, Ch^16^, experimentally derived from endogenous ChIII, is efficiently repaired by homologous recombination (HR), the major DSB repair pathway in fission yeast, using ChIII as a repair template (figs. S1A and S1B) (*10*). In contrast, failed HR repair leads to minichromosome loss (Ch^16^ loss) or to extensive loss of heterozygosity (LOH) predominantly through isochromosome (Ch^I^) formation. Ch^I^ formation results from extensive 5’ end processing of an unrepaired DSB leading to removal of the broken chromosome arm and to replication of the intact arm from inverted repeats within the centromere (*11*). In HR mutants, Ch^I^ formation is significantly increased, suggesting extensive resection associated with failed HR triggers such events (*11*).

To study the kinetics of DSB-induced Ch^I^ formation, we performed a time course using HO endonuclease which generates a DSB uniquely at the *MATa* site introduced within Ch^16^–DSB in a *rad51Δ* background to promote Ch^I^ formation (**Fig. 1A** and **B**). We found that efficient Ch^I^ formation was maximally observed at 96h following HO endonuclease derepression. To test whether the timing of Ch^I^ formation was dependent on the resection distance from the *MATa* break site to the centromere inverted repeats, we moved the *MATa* break site to 10Kb away from the centromere and found Ch^I^ formation was observed much earlier (48h) (**Fig. 1A** and **B**). This suggested that the timing of Ch^I^ formation is proportional to the resection distance from the unrepaired break site to the centromeric inverted repeats.

**Fig. 1.**
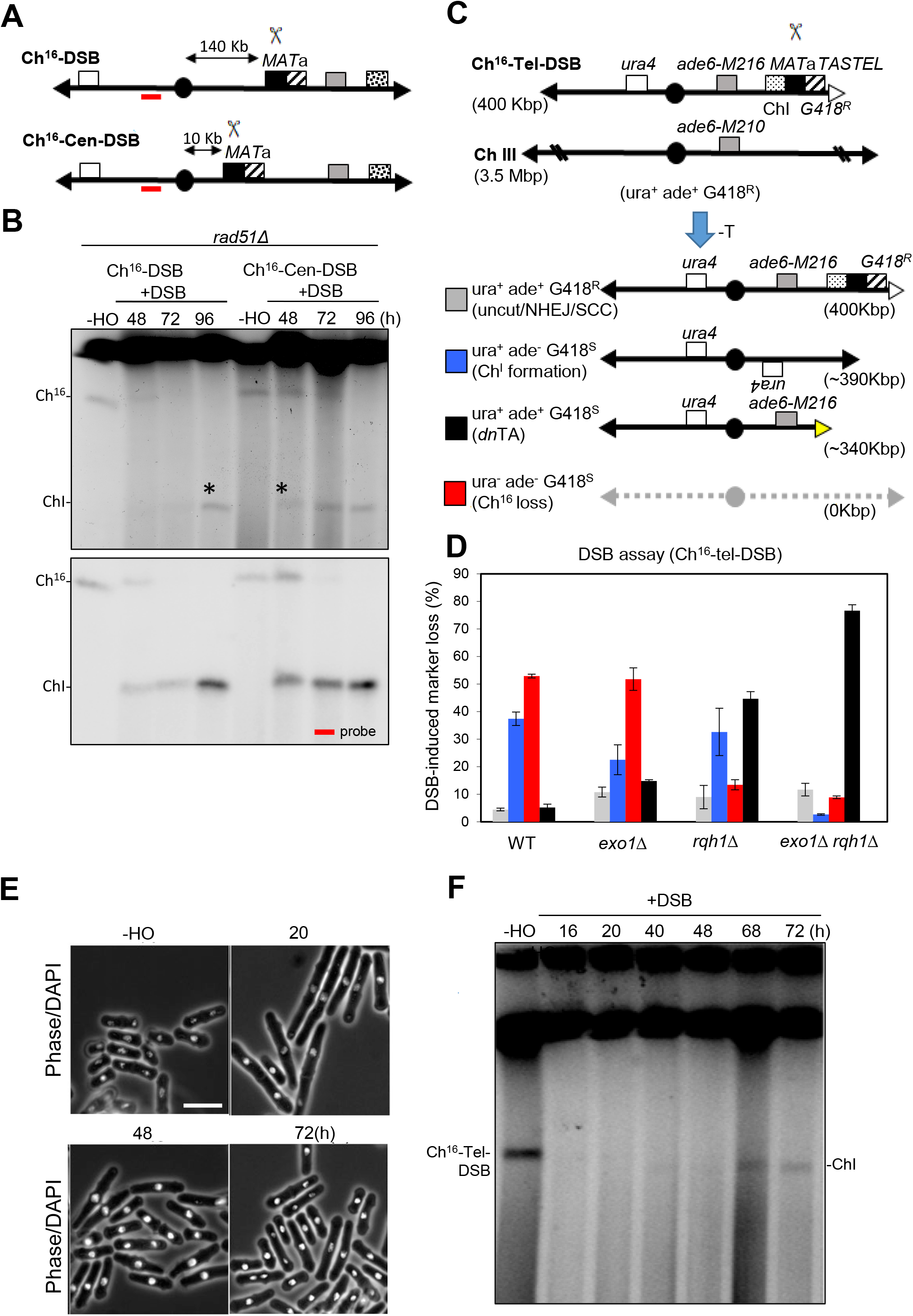
Stabilization of an unrepaired broken chromosome takes multiple generations. **(A)** Schematic of Ch^16^-DSB (Ch^16^-RMGAH) previously described (*11*), in which the distance from *MATa* site to the centromere is 140 Kb; and Ch^16^-Cen-DSB (Ch^16^-R-*mid1*-MGAH) in which the distance from the *MATa* site to the centromere is 10Kb (MATa KanMX6 cassette integrated into the Ch^16^-DSB *mid1* gene). The location of the Ch^16^-DSB probe is indicated (red stripe). **(B)** PFGE analysis of samples taken from *rad51Δ* cells carrying Ch^16^-DSB or Ch^16^-Cen-DSB grown in absence of thiamine (-T) for the indicated times. Bands corresponding to minichromosome (Ch^16^), and isochromosome (Ch^I^) are indicated. Asterisks indicate isochromosome formation. Southern blot analysis of PFGE with probe indicated. **(C)** Schematic of minichromosome Ch^16^-Tel-DSB (Ch^16^-TASTEL) and homologous ChIII. The loci of telomeres (black triangles), the *ura4* gene integrated into the Ch^16^-Tel-DSB *mad3+* locus (white box), centromere (black ovals), *ade6-M216* and *ade6-M210* heteroalleles (gray boxes), 3kb region of homology to ChI (light gray), *MAT*a target site (black box), *KanMX6* gene (G418) (hatched), and artificial telomeric sequence (TASTEL) (white triangle) are indicated. Derepression of *nmt41x-HO* in the absence of thiamine (-T) generates a DSB uniquely at the *MAT*a target site (scissors). Repair of HO-induced DSB by NHEJ or sister chromatid conversion (SCC) if only one sister chromatid is broken) results in retention of all markers resulting in an ura^+^ ade^+^ G418^R^ phenotype, which is indistinguishable from uncut minichromosome. Extensive LOH in which genetic material centromere-distal to the break-site is lost results in ura^+^ ade^-^ G418^S^ phenotype, and results predominantly from isochromosome formation, indicated. Extensive LOH resulting in ura^+^ ade^+^ G418^S^ phenotype usually results from *de novo* telomere addition (dnTA) which can occur between *ade6-M216* and the MATa break site (yellow triangle). Failed DSB repair can also result in loss of the minichromosome, resulting in a ura^-^ ade^-^ G418^S^ phenotype. **(D)** Percentage of DSB-induced marker loss in wild type, *exo1Δ, rqh1Δ*, or *rqh1Δ exo1Δ* backgrounds carrying Ch^16^-Tel-DSB. The levels of uncut/NHEJ/SCC, Ch^16^ loss and LOH are shown. S.E.M values are indicated. The data presented are from at least two independent biological repeats. **(E)** Cell morphology analysis of Ch^16^-Tel-DSB cells grown in absence of thiamine for the indicated times (see also F). Samples were taken at indicated points in parallel to (G) for microscopy analysis. Scale bar = 10 µm. **(F)** PFGE analysis of samples taken from Ch^16^-Tel-DSB cells grown in absence of thiamine for the indicated times (see E) Bands corresponding to minichromosome (Ch^16^-Tel-DSB), and isochromosome (Ch^I^) are indicated.

We considered whether disrupting HR structurally might also generate efficient Ch^I^ formation in a wild-type background. We therefore replaced the Ch^16^ minichromosome arm distal to the *MATa* break site with a synthetic telomere Ch^16^-Tel-DSB thereby abrogating second-end capture (**Fig. 1C**). As predicted, DSB induction and telomere loss in Ch^16^-Tel-DSB in a wild-type background resulted in very high levels of Ch^16^ loss and Ch^I^ formation (**Fig. 1D)**. Extensive LOH was significantly reduced in *rqh1Δ exo1Δ* Ch^16^-Tel-DSB cells, consistent with Ch^I^ formation requiring extensive processing (*11, 12*) (**Fig. 1D**), with LOH resulting instead from *de novo* telomere addition (*dn*TA) (*13*).

A time course following DSB induction in Ch^16^-Tel-DSB indicated Ch^I^ formation was observed at 68h. This timing is in agreement with the expected time to resect 130 kb from the *MATa* break site to the centromere at a resection speed of 4.4 kb/h (*14*). As the length of a normal cell cycle for *S. pombe* is 3.5 h, (*15*), this raised the question as to whether cells continued to divide during this time. Concomitant analysis of cells during the time course analysis indicated that checkpoint-dependent cell elongation was observed at 20h after HO derepression (**Fig. 1E** and fig.S1C and D). However, cells returned to normal length and were observed to proliferate at 48h (**Fig. 1E**), prior to Ch^I^ formation (68h) (**Fig. 1F**). Together, these results strongly suggest that failed HR through telomere loss leads to DNA damage checkpoint adaptation and cell division in the presence of an unrepaired broken chromosome.

Consistent with these observations, deletion of orthologues of known *S. cerevisiae* DNA damage checkpoint adaptation factors (*rad51*Δ, *ku70*Δ, *rdh54*Δ, *rif1*Δ, *srs2*Δ) (*4, 6, 16, 17*) and *rad57*Δ (identified from genome-wide screen in this study) in fission yeast resulted in increased cell death in these mutants following DSB induction within the non-essential Ch^16^-Tel-DSB (**Fig. 2A** and fig.S2A). In contrast, abrogating the DNA damage checkpoint by deleting *rad3+* (ATR) (*18*) in this context did not cause viability loss following DSB induction (fig. S2B). DSB induction in a wild-type strain containing Ch^16^-Tel-DSB was associated with increased Chk1 phosphorylation levels at earlier time points when cell division was arrested, consistent with G2-M checkpoint activation (*19*) (**Fig. 2B**, 20-22h and **Fig. 2C**, 20h). However, Chk1 protein levels were strikingly absent at 25h when cells were observed to initiate cell division (**Fig. 2B**, 25-29h and **Fig. 2C**, 25h).

**Fig. 2.**
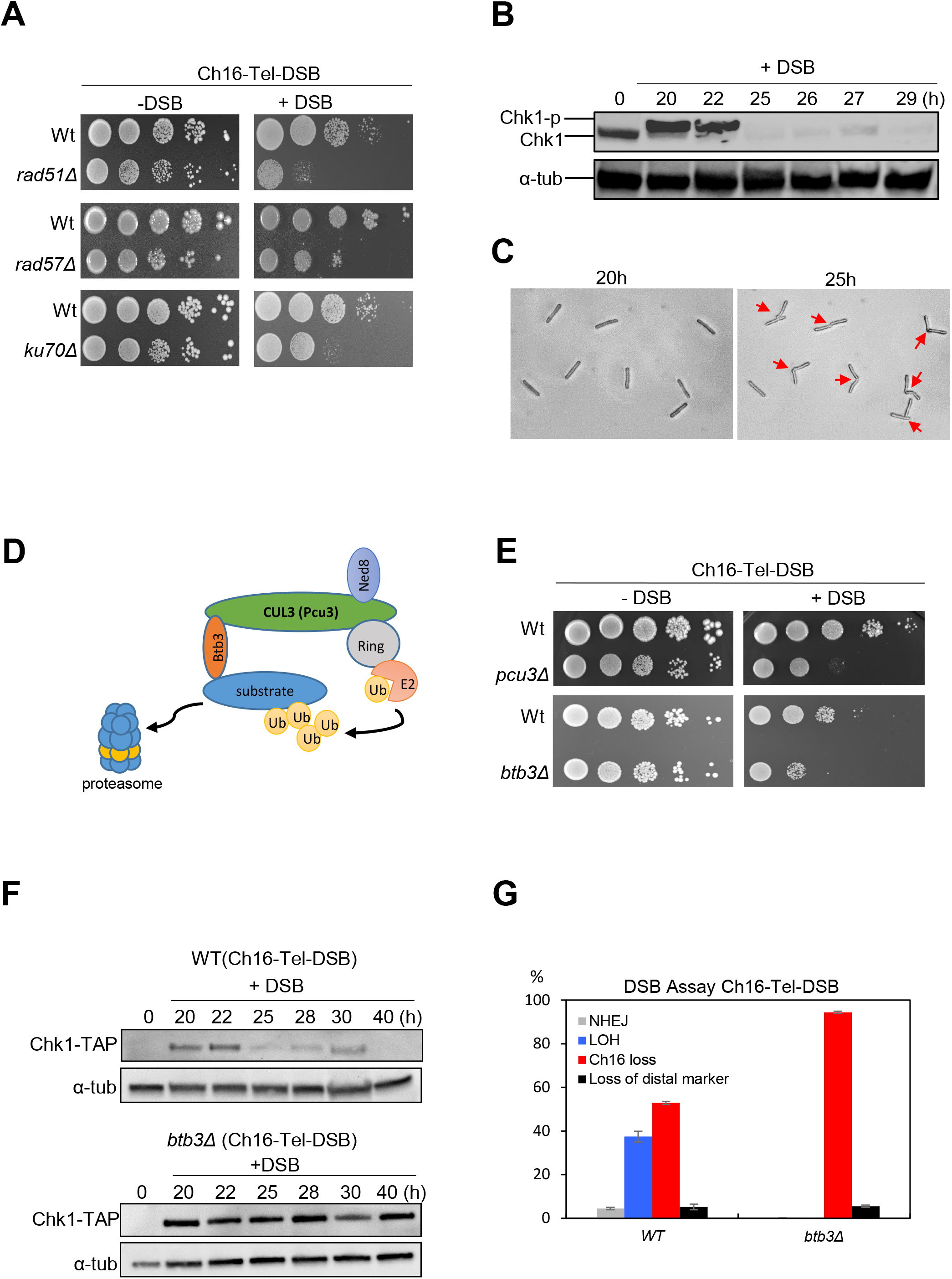
An unrepaired broken chromosomes facilitates checkpoint adaptation through Chk1 loss. **(A)**Viability analysis of wild-type, *rad51*Δ, *rad57*Δ or *ku70*Δ cells carrying Ch^16^-Tel-DSB and *nmt1(41X)-HO* integrated into the *ars1* locus (Williams et al., 2009). Cells were serially diluted (10-fold) and spotted onto Edinburgh Minimal Medium (EMM) + U+A+L+R in the presence (-DSB) or absence of thiamine (+DSB) and incubated at 32°C for 2-3 days. **(B)**Analysis of Chk1 protein levels in cells carrying Ch^16^-Tel-DSB following DSB induction at times indicated. Cells were incubated at 32°C and samples taken at the indicated times following thiamine removal (0h) to derepress *nmt1(41x)-* HO integrated into the *ars1* locus (+DSB) Cell extracts were made by using the TCA method. Tap-tagged Chk1 was detected using an anti-PAP antibody. α-tubulin is shown as a loading control. **(C)**Microscopy analysis of cells taken at 20 hrs and 25 hrs following thiamine removal. Red arrows indicate dividing cells. **(D)**Schematic representation of the Cullin 3-ring ubiquitin ligase (*37*) (upper panel). **(E)**Viability analysis of wild-type, *pcu3*Δ or *btb3*Δ cells carrying Ch^16^-Tel-DSB. Cells were serially diluted (10-fold) and spotted onto EMM plates in the presence (-DSB) or absence of thiamine (+DSB) and incubated at 32°C for 2-3 days (lower panel). **(F)**Analysis of Chk1 protein levels in *pcu3*Δ Ch^16^-Tel-DSB cells following DSB induction at times indicated. Cell extracts were made by using the TCA method (Materials and Methods). Tap-tagged Chk1 was detected using an anti-TAP antibody (upper panel). α-tubulin is shown as a loading control (lower panel). **(G)**Percentage of DSB-induced marker loss in wild type or *btb3Δ* backgrounds carrying Ch^16^-Tel-DSB. The levels of ura^+^ ade^+^ Hyg^R^ (uncut/NHEJ/SCC), ura^-^ ade^-^ Hyg^S^ (Ch^16^ loss), ura^+^ ade^-^ Hyg^S^ (Ch^I^ formation), ura^+^ ade^+^ Hyg^S^ (dnTA) are shown. S.E.M values are indicated. The data presented are from at least two independent biological repeats.

To determine the mechanism of Chk1 loss, we performed a genome-wide screen and found the adaptor protein Btb3 and Pcu3 of the Cullin3/Pcu3 E3 ubiquitin ligase to be required for cell viability of strains carrying Ch^16^-Tel-DSB following DSB induction (**Figs. 2D** and **2E**). Western blot analysis showed that Chk1 protein levels remained high in the absence of Btb3 (**Fig. 2F)**. Pcu3, another component of the Cullin3 complex, was also found to be required for viability following DSB induction, further supporting a role for this complex being required for checkpoint adaptation (**Fig. 2E)**. Moreover, the majority of survivors failed to undergo extensive LOH, and instead had lost Ch^16^-Tel-DSB following DSB induction in a *btb3*Δ background (**Fig. 2G**). These results suggest that the unrepaired broken chromosome leads to Cullin-3-dependent Chk1 loss and checkpoint adaptation.

To explore the fate of the unrepaired broken chromosomes following DNA damage checkpoint adaptation, a single cell microscopy time course was performed to visualise the broken unrepairable Ch^16^-Tel-DSB using Rad52-GFP (*20*). Following DSB induction, Rad52-GFP foci were observed in both daughter cells, consistent with unrepaired broken chromosomes being inherited by both daughters (Movie S1). Surprisingly, Rad52-GFP foci were also observed in both daughter cells for at least two generations following DSB induction (**Fig. 3A**, 180-340 min and Fig. S3, A, B and C). This suggested that the inherited broken minichromosomes were being replicated and segregated in daughter cells. To provide further evidence for replication of inherited broken minichromosomes, Ch^16^-Tel-DSB was visualised following integration of *lacO* repeat arrays into Ch^16^-Tel-DSB and expressing LacI-GFP, which specifically binds to *lacO* arrays. Following break induction within Ch^16^-Tel-DSB, *lacO*/LacI-GFP foci were observed in daughter cells for several generations, consistent with the inherited broken chromosome being efficiently replicated and segregated in daughter cells (**Fig. 3B** and Movie S2).

**Fig. 3.**
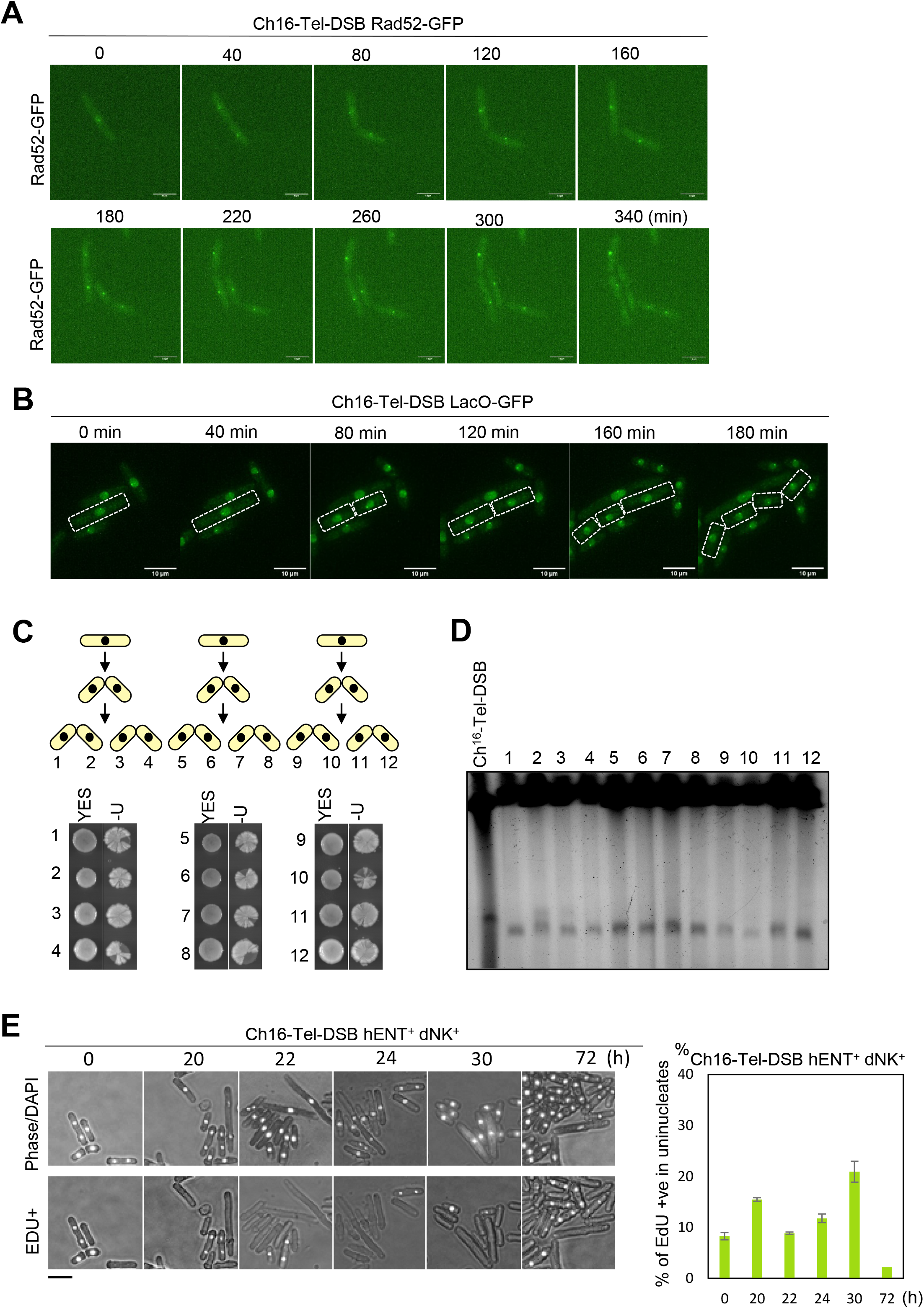
Inherited unrepaired broken chromosomes undergo DNA replication. **(A)**Analysis of Rad52-GFP foci using live-cell imaging of a Ch^16^-Tel-DSB strain encoding Rad52-GFP following DSB induction. Rad52-GFP foci are associated with elongated cells. Rad52-GFP are observed daughter cells over two generations consistent with cell division in the presence of damage, and suggesting that unrepaired Ch^16^-Tel-DSB is segregated and replicated. **(B)** Live-cell imaging of Ch^16^-Tel-DSB lacO strain expressing lacI-GFP following DSB induction. LacO array integrated into the *mid1* gene on the right arm, 10 Kb from the centromere of Ch^16^-Tel-DSB (Materials and Methods). Images show that LacI-GFP foci are observed in each daughter cell for two generations following division of an elongated cell following DSB induction. **(C)** Pedigree analysis of cells carrying Ch^16^-Tel-DSB following DSB induction. Graphic representation of pedigree analysis of single yeast cells and their descendants over two generations (upper panel). An individual elongated cell carrying Ch^16^-Tel-DSB following DSB induction was placed onto a non-selective YES plate and daughter cells separated for two successive generations and incubated to allow colony formation. Colonies were subsequently replica-plated onto EMM-U plates and grown at 32°C for two days. Colony pairs grown on YES and EMM-U plates are shown from three independent analyses (1-4, 5-8, 9-12) (lower panel). **(D)** PFGE analysis of genomic DNA from a wild-type strain containing Ch^16^-Tel-DSB (lane 1) and individual ura^+^ colonies derived from daughter cells from 3C (1-12). **(E)** Analysis of DNA replication in cells carrying Ch^16^-Tel-DSB following DSB induction. Cells were grown in EMM medium following thiamine removal (0hr) and collected at indicated times and treated with 100 μM EdU for 15 min before being fixed in 70% ethanol. Samples were conjugated to Alexa Fluor 488 before being imaged by fluorescence microscopy. DAPI and EdU staining are shown. Quantification of uninucleate cells staining positive for EdU incorporation. S.E.M values are indicated (right panel). Scale bar = 10 µm.

To confirm that unrepaired broken chromosomes are replicated in daughter cells, following DSB induction, individual elongated cells carrying Ch^16^-Tel-DSB were segregated for two generations and allowed to form colonies on non-selective YES plates followed by replica plating onto plates without uracil (ura-). We found all resulting colonies were able to grow on ura-plates from three separate pedigree analyses (**Fig. 3C** colonies 1-12), indicating the unrepaired minichromosome (carrying the *ura4* gene) was inherited in daughter cells. Pulsed Field Gel Electrophoretic (PFGE) analysis showed that all colonies derived from two generations of daughter cells from three independent experiments carried shorter derivatives of Ch^16^-Tel-DSB. These findings confirm that an unrepaired broken chromosome is replicated by successive generations of daughter cells, while undergoing extensive end processing to form Ch^I^ (**Fig. 3D**). Consistent with this, pedigree analysis indicated that the broken minichromosome could be replicated for at least six generations (fig. S3D). Using EdU incorporation, we found that the unrepaired broken chromosome is replicated during normal S-phase and that endogenous DNA replication proceeds normally following adaptation to an unrepaired broken chromosome (**Fig. 3E)**. Following sequence analysis of the left arm of Ch^I^, the number of single nucleotide polymorphisms (SNPs) and indels detected was similar to the wild-type Ch^16^-Tel-DSB strain, consistent with high fidelity DNA replication of the broken chromosome. These results together demonstrate that following checkpoint adaptation, daughter cells inherit an unrepaired broken chromosome, which is subsequently replicated and segregated over multiple generations.

We wished to determine the consequences of such post-adaptive DNA replication and end processing on genome stability in subsequent generations. We noted that colonies derived from single elongated cells carrying an unrepaired broken Ch^16^-Tel-DSB minichromosome when replica-plated from a non-selective (YES) plates to ura-plates exhibited a unique series of colony sectoring patterns of *ura4* marker loss, consistent with break-induced CIN (**Fig. 3C**; fig. S4A). PFGE analysis revealed that CIN was associated with distinct minichromosome sizes in cells from individual LOH colonies (fig. S4B) or from individual cells within the same LOH colony (fig. S4C).

To further examine break-induced CIN across successive generations, pedigree analysis was performed over several generations following DSB induction in Ch^16^-Tel-DSB. We found that 4 of 6 colonies formed from the third generation could grow on ura-plates, consistent with faithful replication of the unrepaired broken Ch^16^-Tel-DSB in successive generations (fig. S4D). Unrepaired broken Ch^16^-Tel-DSB carrying *ura4* were found to give rise to both *ura+* and *ura-* daughters, consistent with Ch^16^ loss. Moreover, *ura-* colonies were found to give rise to *ura+* colonies, suggesting that the parental cell had lost the unstable minichromosome in subsequent divisions.

These findings raised the possibility that independent processing of replicated broken chromosomes by daughter cells might contribute to such widespread CIN. Pedigree analysis of DSB repair outcomes in daughter cells following DSB induction in the parental cell, carrying the repairable minichromosome Ch^16^-DSB, revealed GC/GC (46.2%) GC/ (NHEJ or SCC) (0.8%); GC/LOH (12.0%); GC/Ch^16^ loss (6.7%); LOH/LOH (13.6%); LOH/Ch^16^ loss (11.0%); or Ch^16^ loss/ Ch^16^ loss (2.8%) daughter colony pairs (fig. S4E). Similarly, DSB induction in the parental cell carrying the unrepairable minichromosome Ch^16^-Tel-DSB revealed LOH/LOH (55%), LOH/dead (19%) Ch^16^ loss/LOH (0.5%), dnTA/LOH (6%) dnTA/dnTA (2%) dead/dnTA (1%) and dead/dead (17%) daughter colony pairs (fig. S4F). Thus, broken sister chromatids can be repaired or misrepaired independently, thereby contributing to genetic variation and CIN after each successive division.

Next generation sequencing (NGS) analysis of genomic DNA from LOH strains exhibiting different minichromosome sizes (**Fig. 4A** and fig. S4G) confirmed that the left arm of the minichromosome had been duplicated in LOH1 and 9 consistent with Ch^I^ formation (**Fig. 4B** and figs. S4, H and I). LOH9 exhibited a duplication of a larger portion the centromeric region, explaining the increased minichromosome size compared to LOH1 (fig. S4I). In contrast, LOH5, was found to have retained some of the right arm, consistent with *de novo* telomere addition (*13, 21*) (fig. S4J). Importantly, NGS analysis of these LOH strains also revealed various levels of amplification or deletions of endogenous chromosomes indicating unrepaired broken chromosomal rearrangements can also cause genome-wide chromosome rearrangements (**Fig. 4B** and figs. S4, H, I and J).

**Fig. 4.**
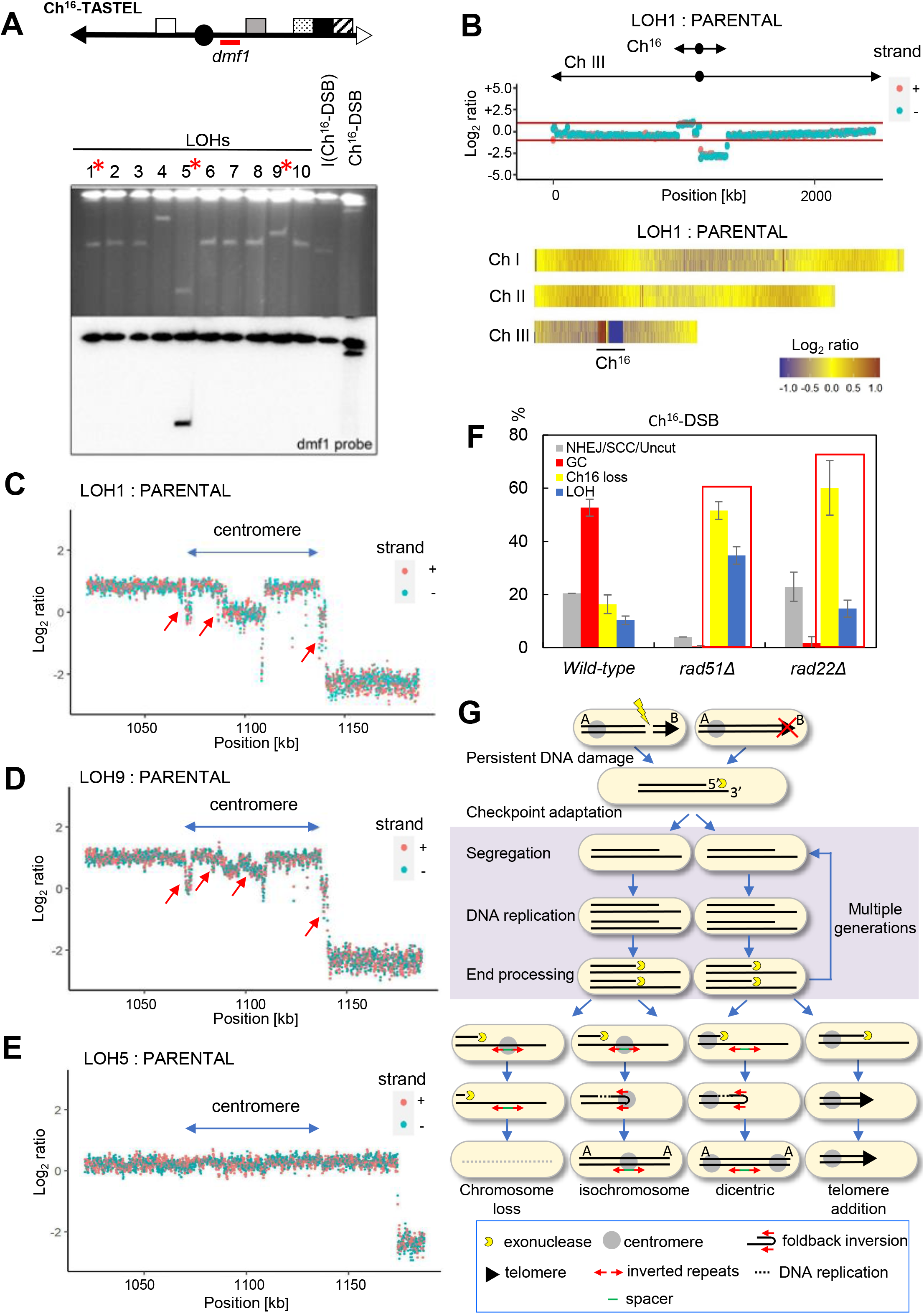
An unrepaired broken chromosome can lead widespread chromosomal instability over multiple generations. **(A)** Schematic indicating *dmf1* probe localization within Ch^16^-Tel-DSB (upper panel). PFGE analysis of chromosomal DNA from wild-type Ch^16^-Tel-DSB and individual wild-type ura^+^ ade^-^ G418^S^ (LOH) strains isolated after DSB induction. Southern blot of the PFGE probed with *dmf1* (lower panel). **(B)** Next generation sequence analysis of LOH strains. NGS analysis showing the log2 of the signal ratio between parental Ch16-Tel-DSB and LOH1 strain across Ch III. Locations of Ch^16^ and Ch III centromeres (oval) and telomeres (arrows) are indicated. Data acquisition and normalization were carried out as described in Materials and Methods. Heat map representation of NGS analysis of LOH1 showing three endogenous chromosomes. Yellow indicates a 1:1 ratio, red indicates signal intensity >1 and blue is <1. **(C)** NGS analysis of LOH1 across centromeric genome sequences. Red arrows point out spacer regions between deleted and duplicated segments. **(D)** NGS analysis of LOH9 across centromeric genome sequences. Red arrows point out spacer regions between deleted and duplicated segments. **(E)** NGS analysis of LOH5 across centromeric genome sequences. **(F)** Percentage of DSB-induced marker loss in a wild-type, *rad51Δ* or *rad52Δ* background carrying Ch^16^-RMGAH. The levels of uncut/NHEJ/SCC, Ch^16^ loss and LOH are shown. S.E.M values are indicated. The data presented are from at least two independent biological repeats. Red boxes indicate misrepair outcomes in HR mutant cells. **(G)** SERPent-cycle induced chromosomal instability model. Persistent DNA damage leads to post-adaptive cycles of segregation, DNA replication and end processing (highlighted in purple). Observed and predicted outcomes are shown.

Further NGS analysis of LOH1 and 9 suggested more complex genome rearrangements within the centromere (**Fig. 4C** and **4D**), in contrast to LOH5 (**Fig. 4E**). ‘‘Spacer’’ regions across centromeric inverted repeats (imr, dg, dh and irc) were present but not amplified (**Fig. 4C** and **4D**). This is consistent with breakpoint resection exposing single-strand inverted repeat regions undergoing intrastrand annealing resulting in formation of a ‘hairpin’ or ‘fold-back’ looping structure which may lead to duplication of the intact left arm and Ch^I^ formation (*22, 23*). While unrepaired breaks can lead to increased Ch^16^ loss and Ch^I^ following Rad51 loss (**Fig. 4F**), Ch^16^ loss was significantly increased while Ch^I^ levels were similar to wild-type following Rad52 loss (**Fig. 4F**). These results are consistent with a role for Rad52 in promoting efficient HR repair (*24*), and the reduced levels of ChI formation in *rad52*Δ compared to *rad51*Δ suggests a further role for Rad52 in facilitating single-strand annealing of resected inverted repeats, thereby triggering chromosome arm duplication and Ch^I^ formation (*11, 23, 25*). Together, the above findings indicate that an unrepaired broken chromosome can lead to widespread CIN over multiple generations.

Here we propose a new model for CIN in which an unrepaired broken chromosome is subject to post-adaptive cycles of <u>se</u>gregation, DNA replication and DNA end <u>p</u>rocessing, (SERPent cycles), in successive daughter cells, thereby driving widespread CIN across the resulting population (**Fig. 4G**). A key feature of SERPent cycle-induced CIN is that following checkpoint adaptation, inherited unrepaired broken chromosomes are replicated in daughter cells. This finding has considerable significance as this explains how genome instability can be spread across the population of daughter cells rather than being otherwise limited to a single cell lineage. Further, replication of unstable intermediates provides a mechanistic basis for rapid genetic variations, thereby acting as an engine to drive further CIN prior to chromosomal stabilization or loss.

Our findings extend our previously proposed model for Ch^I^ formation in which extensive exonucleolytic resection proceeds over many generations. This facilitates resection through large inverted repeats within the Ch^16^ centromere and their annealing to form an intrastrand hairpin loop structure, which promotes DNA replication leading to a large inverted chromosomal duplication and to Ch^I^ formation (*11, 25-27*) (**Fig. 4G**). This model predicts such events will be more frequent when HR repair is abrogated structurally or genetically, thereby leading to extensive resection; that inactivation of the DNA damage checkpoint will also increase such genome instability; the number of SERPent cycles will be proportional to the distance resected from the break site to the inverted repeats, and the type of chromosomal duplication arising will depend on the location of inverted repeats in relation to the centromere, as while annealing of resected inverted repeats within the centromere facilitates Ch^I^ formation, annealing of resected inverted repeats distal to the centromere will generate dicentrics (*23*) (**Fig 4G**). Our findings further predict that SERPent cycles will precede fold-back inversions. Evidence for fold-back inversions has been found in the genomes of viruses (*28*), prokaryotes (*29*), is associated with genetic disease (*22, 30, 31*) and a variety of cancers (*32-34*) Moreover, fold-back inversions observed in human cancer cells having escaped telomere-driven crisis occur independently of DSB repair (*35, 36*), consistent with our findings. Thus, we anticipate that this study will contribute to the understanding of rapid genome evolution, pathogenesis and tumorigenesis.

## Author contributions

C-C.P, S.C.D and T.C.H authored the manuscript and designed experiments. All authors contributed to and commented on the manuscript. Experiments in Fig. 1 were performed by C-C.P; in Fig. 2 were performed by C-C.P; in Fig. 3 were performed by C-C.P and S. C. D; in Fig.4 were performed by C-C.P and W-C.C. Fig. S1 was performed by C-C.P; Fig. S2 by C-C .P and S.C.D.; Fig. S3 by C-C.P.; Fig.S4 by C-C.P., W-C.C and S.C.D.; C. W performed Southern blotting analysis; W-C. C and S. E. K contributed NGS data analyses; C-C.P, N-Y.C and S.C.D performed the Bioneer genome-wide screen in fission yeast and C-C P validated results.

## Data and materials availability

All materials and raw data are available upon request.

## Supplementary Materials

Figs. S1 to S4

References (*38-41*)

Movies S1 and S2

## Materials and Methods

Standard media and growth conditions were used throughout this work. Cultures were grown in rich media (YE6S) or Edinburgh minimal media (EMM) at 32 °C with shaking, unless otherwise stated.

### DNA double strand break assay

Assay in minichromosome Ch^16^-Tel-DSB strain was carried out as described previously (38). The percentage of colonies undergoing NHEJ/SCC (ura^+^ ade^+^ G418^R^), minichromosome Ch^16^ loss (ura^-^ ade^-^ G418^S^) or LOH (ura^+^ ade^-^ G418^S^; ura^+^ ade^+^ G418^S^) were calculated. More than 1000 colonies were scored, and each experiment was carried out independently three times.

### Pulse Field Gel Electrophoresis

The procedures used in this study for PFGE analysis have been described previously (39). For the time course experiment, Ch^16^-Tel-DSB cells were inoculated in EMM+U+A+L+R medium (+T or -T). Samples were collected and washed twice in 0.05 M EDTA at times indicated before PFGE analysis. Southern blots were carried out as previously described by (11)

### Pedigree analysis

The procedures used in this study for pedigree analysis have been described previously (21).

### Protein analysis

Protein extracts were made by TCA extraction and analysed by Western blotting as described previously (40). TAP-tagged proteins were detected with peroxidase–anti-peroxidase– soluble complex (P1291, Sigma). α-tubulin was detected with antibody T5168 (Sigma).

### Microscopy analysis

Ch^16^-Tel-DSB cells were inoculated in EMM medium in the presence or absence of thiamine at 32°C. Samples were collected at indicated time points, fixed in methanol/acetone, rehydrated and stained with 4′,6-diamidino-2-phenylindole (DAPI) before examination using Zeiss Axioplan 2ie microscope, Hamamatsu Orca ER camera. Open source micromanager software was used to analyse the image. For Edu (5-ethynyl-2′-deoxyuridine) incorporation and detection, Ch^16^-Tel-DSB cells were grown in EMM medium in the absence of thiamine for 72 hrs. Samples were collected at indicated time points and treated with 100 μM EdU for 15 min before being fixed in 70% ethanol. The procedures used in this study for Edu detection have been described previously (41)

### Live Cell Imaging

Live cell imaging was performed on agarose pads as previously described (Merlini et al., 2017) with the following modifications; EMM was prepared by filter sterilisation and supplemented with 225mg/l adenine, uracil, lysine, leucine, and arginine with 2% ultrapure agarose. 50µl of cell suspension at approximately 10^6^ cells/ml were placed onto the agarose pad and cover slips were fixed in place with Valap. Imaging was performed on a Nikon Ni-E microscope inverted microscope at 100 X magnification in a temperature-controlled environment chamber at 32°C. Multiple XY positions were taken and two-channel images (transmitted light and GFP) were taken over 5 Z-stacks of 0.4µm. Imaging was performed every 20 minutes for up to 18 hours. Deconvolution of the GFP channel was performed using Nikon’s NIS-Elements software using an automatic algorithm. The images shown are maximum projections of the deconvolved image GFP-channel and a single-plane image from the transmitted light channel.

### Whole genome DNA sequencing

S. pombe DNA was extracted from cells grown to log phase at 32°C using MasterPureTM Yeast DNA purification kit (Lucigen/Epicentre). Genomic DNA of Ch^16^-Tel-DSB, LOH1, LOH5 and LOH9 was send to Novogen (UK) Company Limited for whole genome sequencing (Illumina PE150). The resulting reads were aligned to the reference genome Schizosaccharomyces pombe ASM294v2 using bowtie2 v2.2.6. Only the locations with best alignment score are kept if the reads are aligned to multiple locations. The average coverage rate over all samples and all chromosomes is 64.7 reads. The number of reads over equally sized regions are counted by python package HTSeq.

**Fig. S1.**
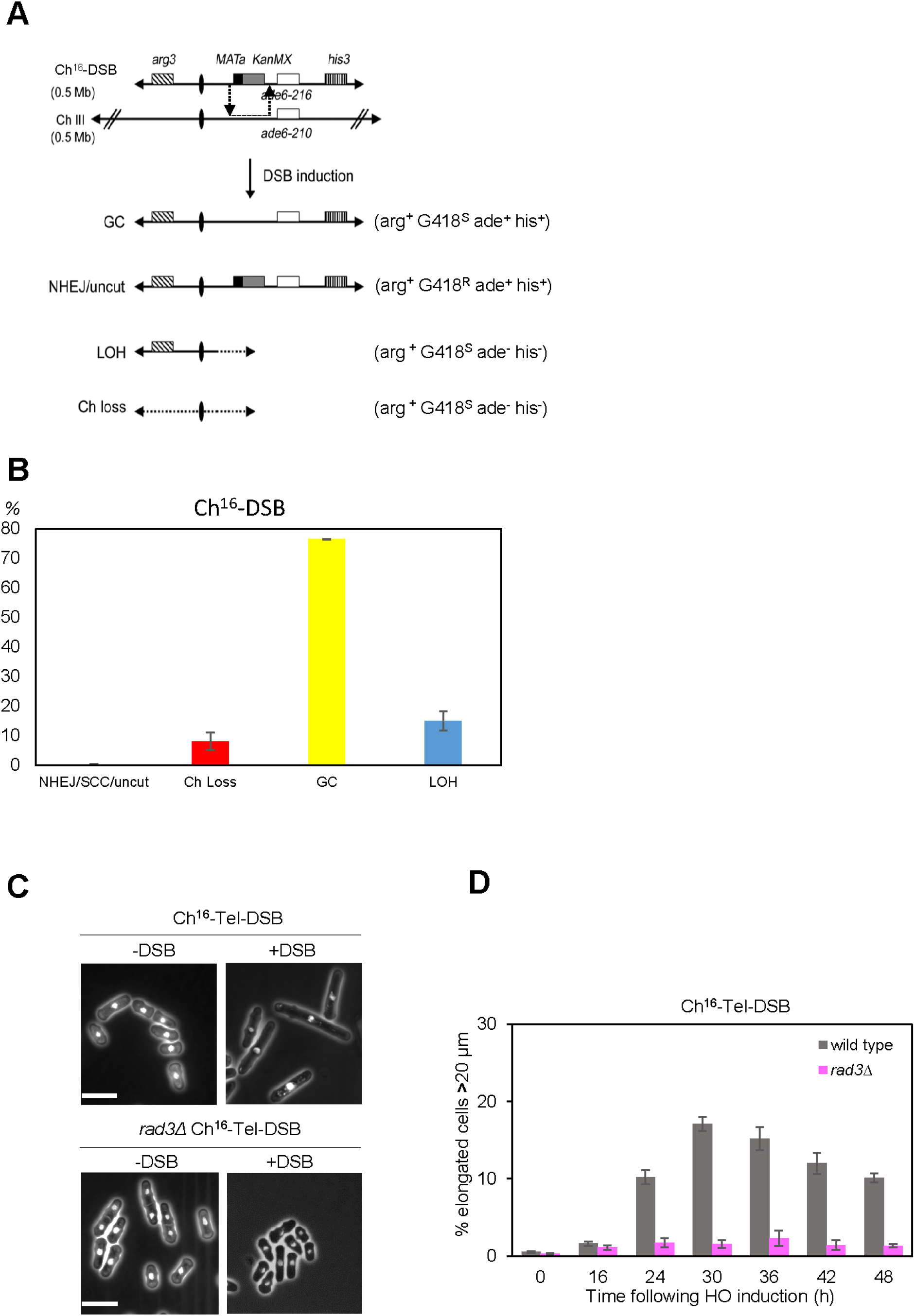
DSB repair outcomes following a site specific DSB induction in Ch^16^-DSB.(**A**) Schematic of Ch^16^-DSB. Ch^16^-DSB, ChIII, centromeric regions (ovals), complementary heteroalleles (*ade6-M216* and *ade6-M210*; white), and the *his3* marker (white) was inserted ∼50kb downstream from *ade6-M216*, as previously shown (11). The *MATa* site (black) with an adjacent *kanMX6* resistance marker (grey) was inserted into *spcc23B6*.*06* ∼ 30kb upstream from *ade6-M216*. The *arg3* marker was inserted into *spcc1795*.*09* on the left arm of the minichromosome. Derepression of pREP81X-HO (not shown) generates a DSB uniquely at the *MATa* target site (scissors). Repair of HO-induced DSB by NHEJ results in retention of all markers resulting in an arg^+^ G418^R^ ade^+^ his^+^ phenotype as indicated. DSB repair by sister chromatid conversion (SCC) during S or G2 phase, in which one of the two sister chromatids is intact, and used as a repair template results in retention of all markers, resulting in an arg^+^ G418^R^ ade^+^ his^+^ phenotype, as indicated. This is indistinguishable from NHEJ in a wild-type background. DSB repair by interchromosomal gene conversion (GC) results in loss of the *KanMX* gene adjacent to the *MATa* break site while the other markers are retained resulting in an arg^+^ G418^S^ ade^+^ his^+^ phenotype. Failed DSB repair results in loss of the minichromosome and loss of all of the markers, resulting in an arg^-^ G418^S^ ade^-^ his^-^ phenotype, as indicated. Extensive loss of heterozygosity (LOH) in which genetic material centromere-distal to the break-site is lost results in an arg^+^ G418^S^ ade^-^ his^-^ phenotype, as indicated. LOH can result from cross-overs associated with gene conversion, break-induced replication, de novo telomere addition or isochromosome formation (11; 21). (**B**) Percentage of DSB-induced marker loss in wild-type cells containing Ch^16^-DSB. The levels of non-homologous end joining/sister chromatid conversion/uncut (NHEJ/SCC/uncut), gene conversion (GC), minichromosome loss (Ch^16^ loss), and LOH are shown. s.e.m. values are indicated. The data presented are from at least two independent biological repeats. (**C**) Cell morphology analysis of Ch^16^-Tel-DSB *or rad3Δ* Ch^16^-Tel-DSB cells following DSB induction. Prior to DSB induction (-DSB), all cells are normal length (12µm). Upon DSB induction (+DSB), *Ch*^*16*^*-Tel-DSB* cells undergo cell cycle arrest, resulting in an elongated phenotype (>20mm) after 20h. In contract, *rad3Δ Ch*^*16*^*-Tel-DSB* cells do not show elongated phenotypes. Size bar = 10µm (**D**) Percentage of cell population with elongated phenotype (>20µm) at time points indicated following *pREP81x-HO* derepression following thiamine removal (+DSB) in wild-type or *rad3*Δ cells carrying *Ch*^*16*^*-Tel-DSB*.

**Fig. S2.**
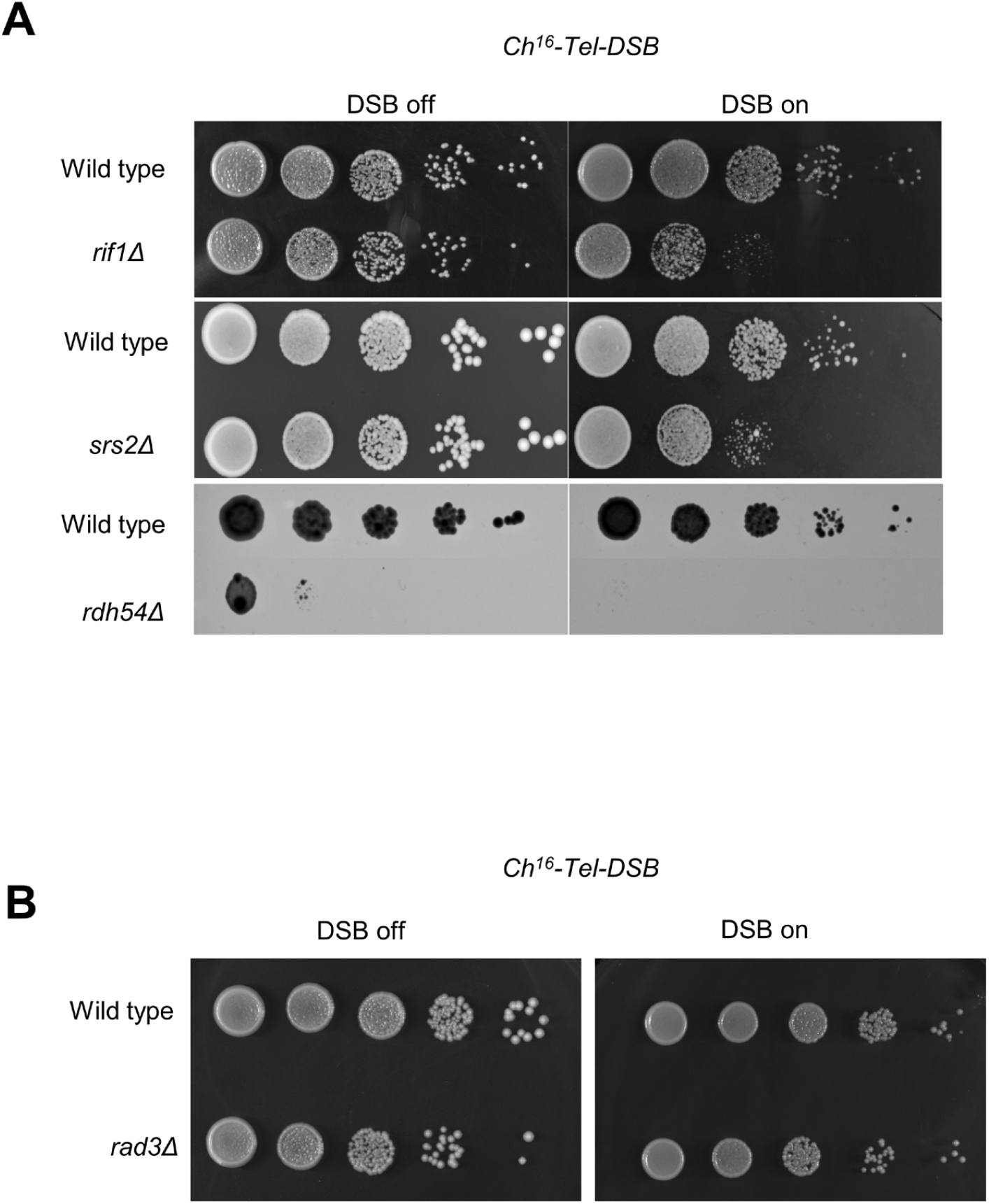
Cell division is facilitated by DNA damage checkpoint adaptation. (**A**) Serial dilution of *Ch*^*16*^*-Tel-DSB, srs2Δ Ch*^*16*^*-Tel-DSB, rif1Δ Ch*^*16*^*-Tel-DSB* and *rdh54Δ Ch*^*16*^*-Tel-DSB* strains in the presence or absence of thiamine. Plates were incubated at 32°C for 3 days. (**B**) Serial dilution of *Ch*^*16*^*-Tel-DSB, rad3Δ Ch*^*16*^*-Tel-DSB* strains in the presence or absence of thiamine. Plates were incubated at 32°C for 3 days.

**Fig. S3.**
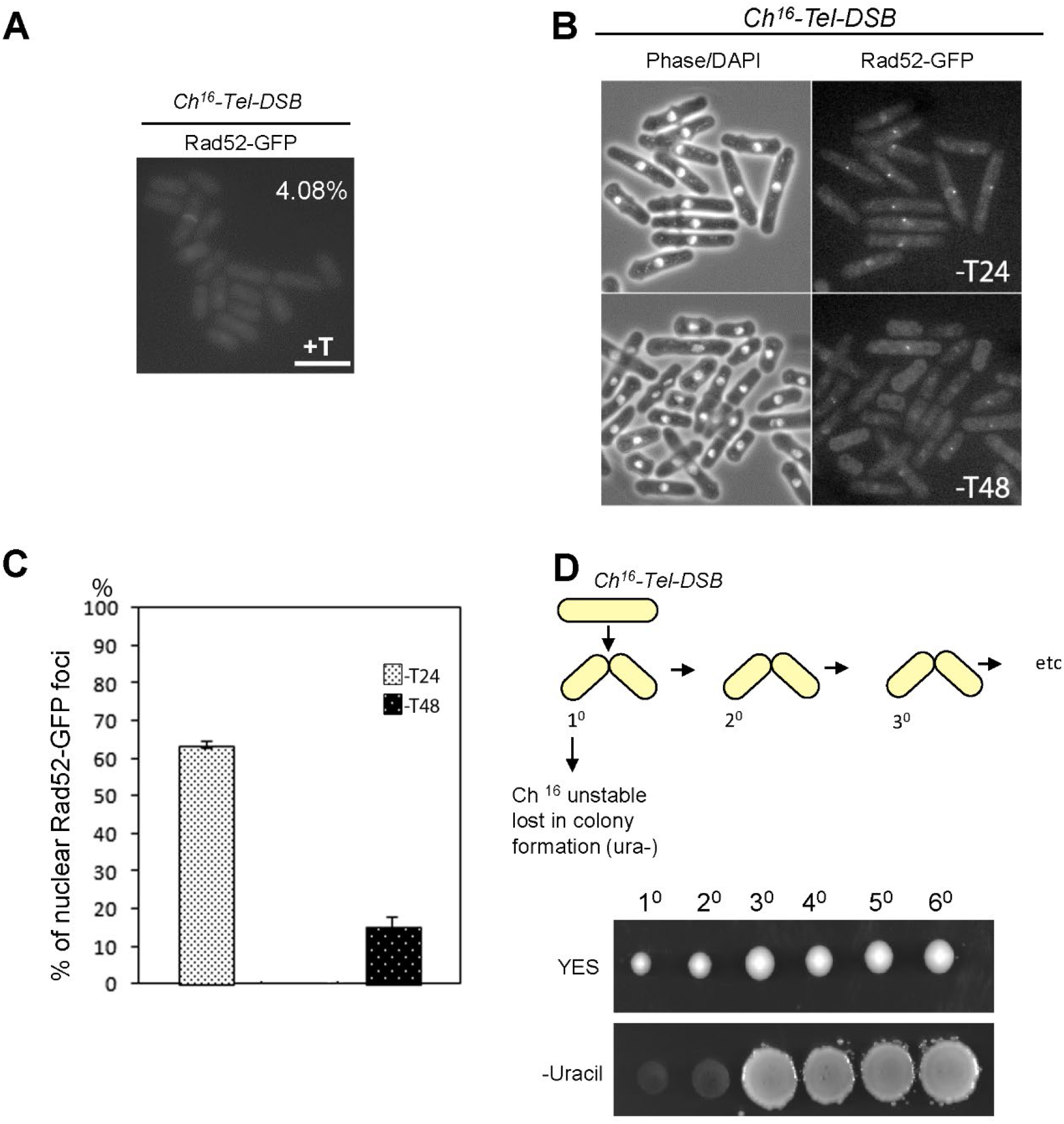
Analysis of Rad52 foci in *Ch*^*16*^*-Tel-DSB* cells following DSB induction. (**A**) Microscopic analysis of Rad52-GFP foci in *Ch*^*16*^*-Tel-DSB* cells in the absence of DSB induction; + T indicates in the present of 5 µg/ml thiamine. Scale bar =10 µm. (**B**) Analysis of Rad52-GFP foci in *Ch*^*16*^*-Tel-DSB* cells at 24 hrs and 48 hrs following DSB induction. -T indicates in the absence of thiamine. (**C**) Quantification of Rad52-GFP foci in *Ch*^*16*^*-Tel-DSB* cells following DSB at indicated time points in B. Data are the mean of two experiments and error bars (±s.e.) are shown. (**D**) Representative pedigree of 6 sequential daughters from a single elongated *Ch*^*16*^*-Tel-DSB* cell following DSB induction. 1° signifies the colony arising from the first daughter cell, 2° signifies the colony arsing from the second, 3° signifies the colony arising from the third, 4° signifies the colony arising from the forth daughter cell, 5° signifies the colony arising from the fifth daughter cell, and 6° signifies the colony arising from the six daughter cell. These sequential daughter cells were then replica plated onto uracil-plates to show the presence or absence of minichromosomes.

**Fig. S4.**
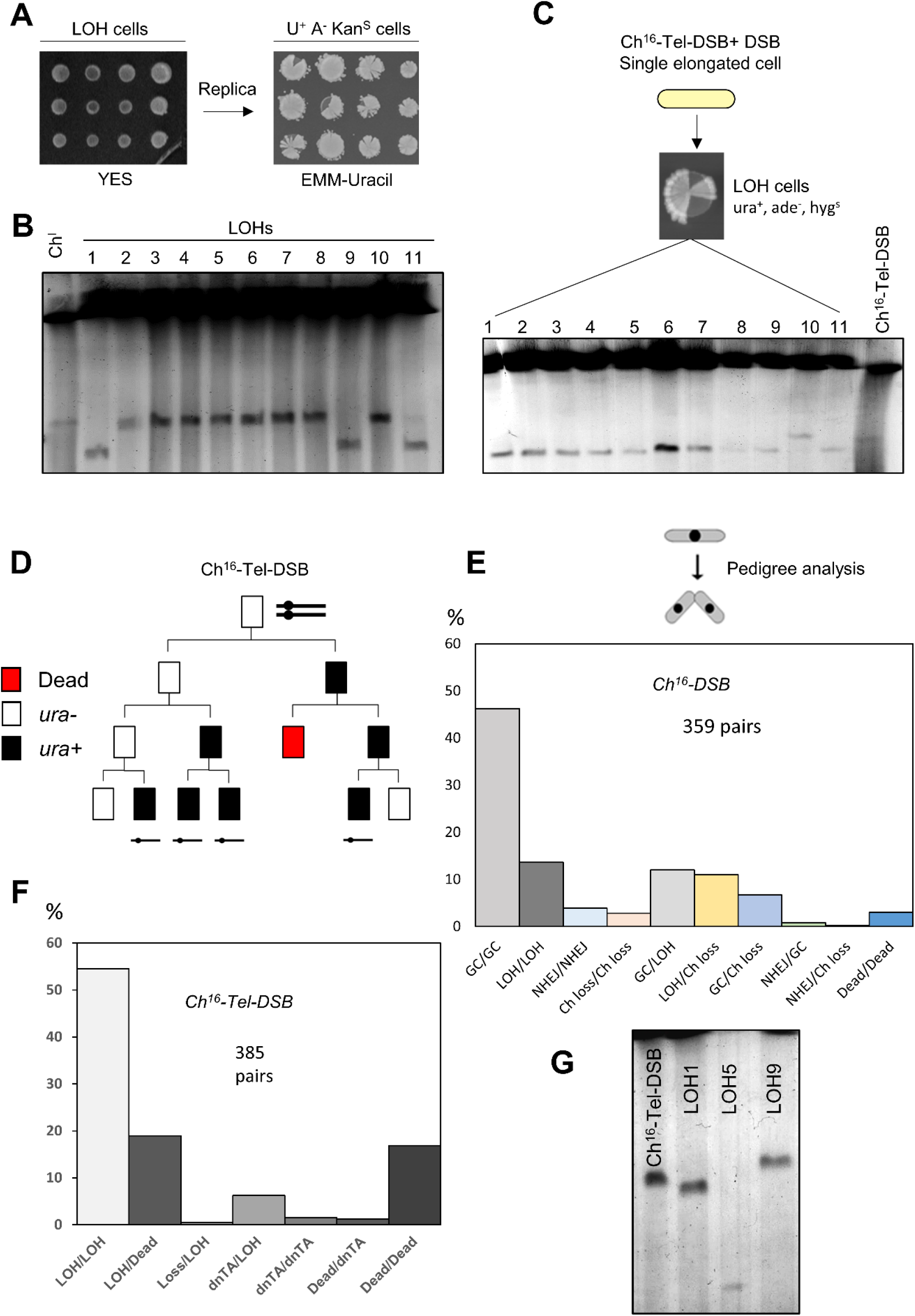
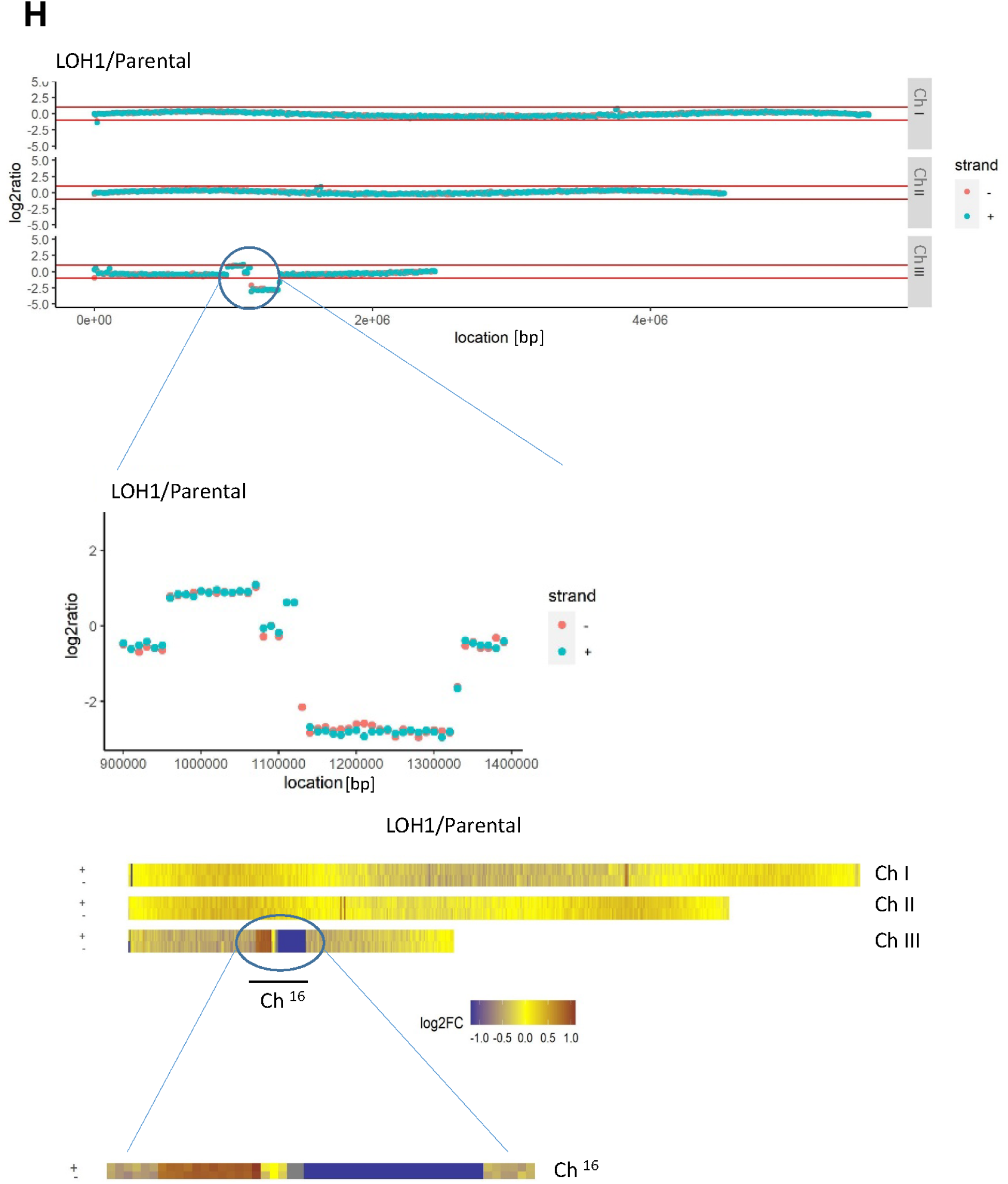
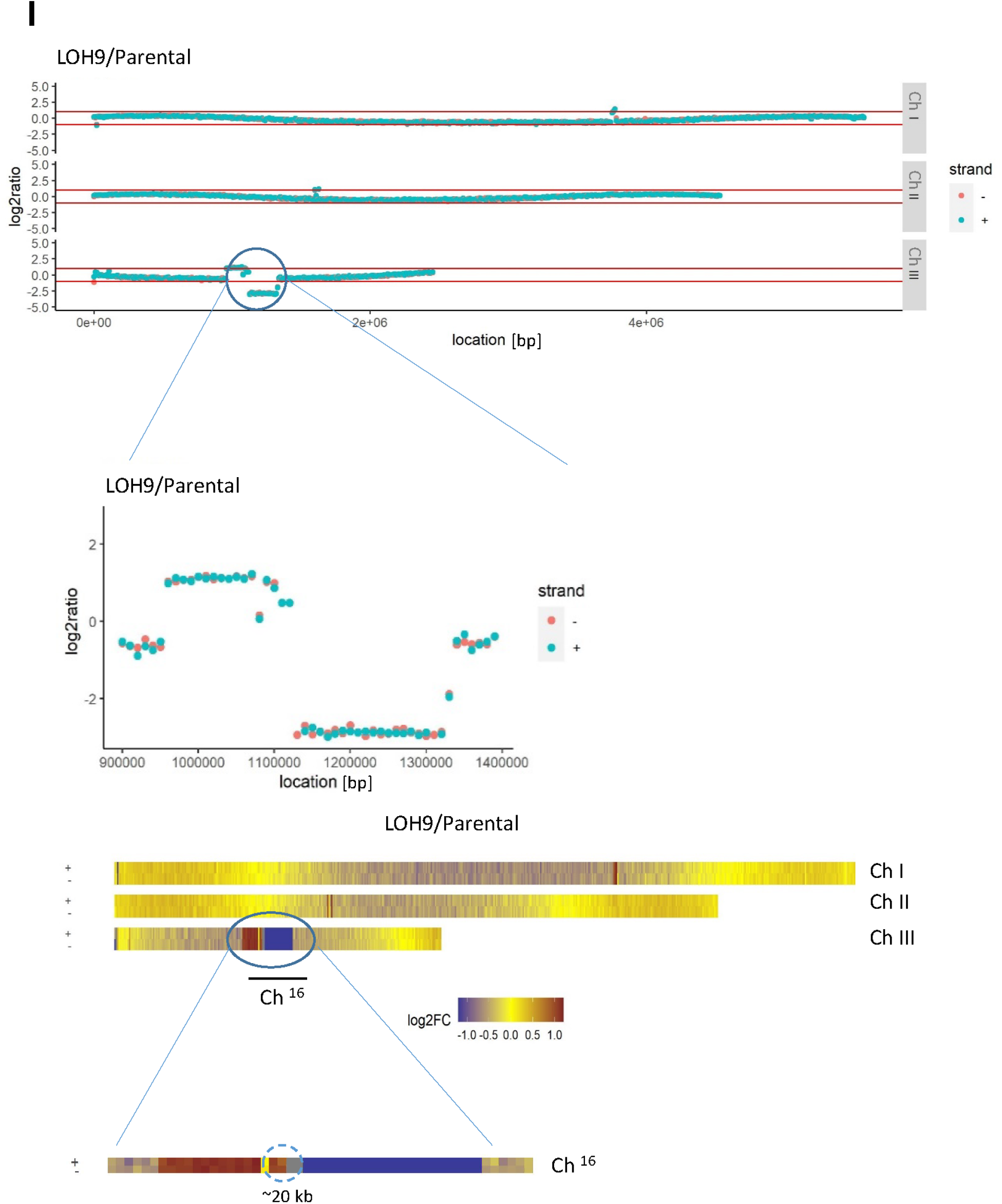
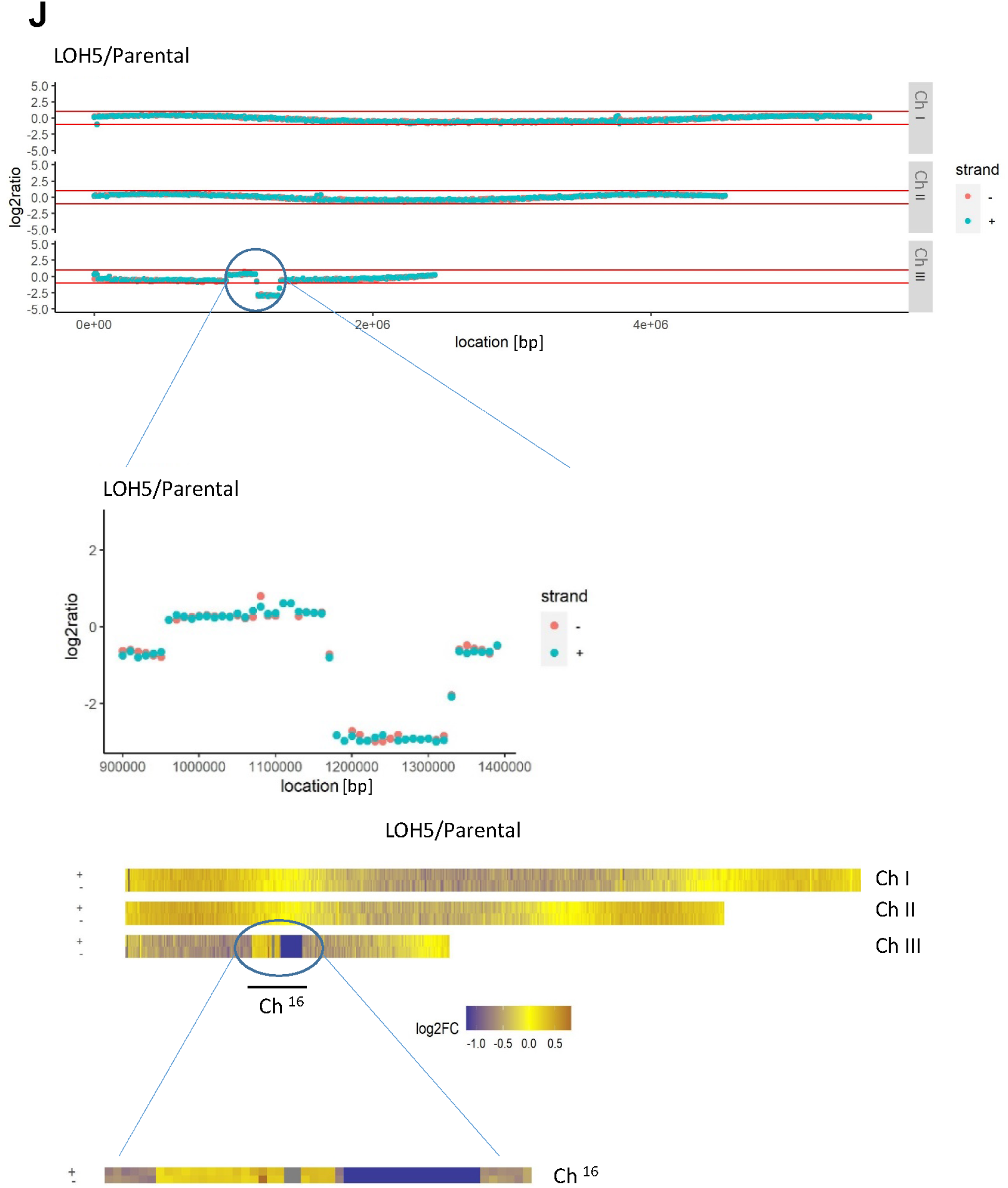
A single DSB can lead to a spectrum of chromosomal rearrangements over generations. (**A**) Sectoring analysis of colonies formed from individual elongated Ch^16^-Tel-DSB cells grown on YES plates following DSB induction. Unselected colonies were replica plated onto uracil-plates and incubated at 32°C for 2-3 days. (**B**) PFGE analysis of chromosomal DNA from sectoring colonies described in (A) (lower panel). (**C**) PFGE analysis of chromosomal DNA from wild type Ch^16^-Tel-DSB descendants (1-11) derived from a single elongated Ch^16^-Tel-DSB cell following DSB induction. (**D**) Pedigree analysis of dividing Ch^16^-Tel-DSB cells following DSB induction. Daughter cells were separated over several cell divisions and were allowed to form colonies on unselective plates. Colonies were replica plated and scored for growth on uracil-plates to identify the presence or absence of minichromosomes. (**E**) DSB repair outcomes in paired daughter cells following a site specific DSB induction in Ch^16^-DSB. (**F**) DSB repair outcomes in paired daughter cells following a site specific DSB induction in Ch^16^-Tel-DSB. (**G**) High resolution PFGE analysis from wild-type Ch^16^-Tel-DSB, individual wild-type ura^+^ ade ^+^ G418^S^ (LOH1, LOH5 and LOH9) strains isolated after DSB induction. (**H**) NGS analysis of LOH1 in (G). NGS analysis showing the log2 of the signal ratio between parental Ch^16^-Tel-DSB and LOH1 strain. Data acquisition and normalization were carried out as described in Materials and Methods. LOH1/Parental density heat-map displayed on three endogenous chromosomes. Yellow indicates a 1:1 ratio, the red is higher than 1 and blue is lower than 1 (**I**) Similar analysis was carried out for LOH9 compared to parental CH^16^-Tel-DSB. (**J**) Similar analysis was carried out for LOH5 compared to parental CH^16^-Tel-DSB

**Movie S1**.

Live cell imaging of CH^16^-Tel-DSB with rad22-GFP after HO-induction. Rad22 foci are seen in the first elongated cell, indicating that rad22 associates with the HO-induced DSB. The foci are still present in subsequent daughter and granddaughter cells suggesting that the DSB is inherited into the next generation. Details for live cell imaging microscopy are described above.

**Movie S2**.

Live cell imaging of CH^16^-Tel-DSB containing a lacO array and lacI-GFP after HO-induction. LacI-GFP foci are seen in the first elongated cell, showing the presence of Ch^16^ in a DNA damage checkpoint activated cell, suggesting the presence of a HO-induced DSB. The lacI-GFP foci are still present in subsequent daughter and granddaughter cells suggesting that Ch^16^ is replicated in future generations. Details for live cell imaging microscopy are described above.

